# Memory-Related Default-Executive Coupling Across the Lifespan and Associations with Changes in Cognitive Control

**DOI:** 10.1101/2025.09.02.673776

**Authors:** Eirik G. Blakstad, Markus H. Sneve, Didac Vidal-Pineiro, Kristine B. Walhovd, Anders M. Fjell, Håkon Grydeland

## Abstract

Episodic memory and cognitive control declines in aging. The default-executive coupling hypothesis of aging (DECHA) suggests that a neural correlate of cognitive decline in aging is increased functional connectivity (FC) between lateral prefrontal areas and the default mode network. Here, in a lifespan sample (n=552, 6-81years), we tested FC between the left dorsolateral prefrontal cortex (dLPFC) and the default mode network (DMN) during the encoding and retrieval phases of an episodic memory fMRI task. We created two age groups based on evidence for episodic memory decline after 30 years: a youth group encompassing childhood, adolescence, and young adulthood (6-29 years) and an aging group (30-81 years). To test if the dLPFC-DMN FC was associated with changes in cognitive control, we used longitudinal change (up to 10 years) in a Stroop inhibition/switching task linked to the prefrontal cortex. Results showed (i) lower dLPFC and DMN connectivity with age in the youth group and higher connectivity in the aging group, and (ii) this FC was associated with an age-related increase in the inhibition task completion time, particularly in the aging group. However, dLPFC showed similar relationships with other networks, particularly salient attentional and control subnetworks, and despite decline in cognitive control associated with memory performance, memory-related FC between dLPFC and DMN did not. Although this link with memory performance remains unclear, the results using longitudinal cognitive data align with the DECHA mechanisms and extends the current proposal by indicating inverse relationships in development and the relevance of additional attentional and control network coupling.

## 1 Introduction

In aging, while a few cognitive processes like semantic knowledge are relatively resistant, most cognitive processes deteriorate, including episodic memory and inhibitory control functions (Park & Reuter-Lorenz, 2009). This age-related decline in cognitive processing is accompanied by an age-related shift in structural and functional brain properties (Chadick et al., 2014; Turner & Spreng, 2015). To account for these observations, Spreng and Turner (2019) proposed the default-executive coupling hypothesis of aging (DECHA) (Spreng & Turner, 2019; Turner & Spreng, 2015). DECHA posits that cognitive decline is accompanied by an increased and inflexible coupling between the default-mode network (DMN) and lateral prefrontal regions paralleling decline in cognitive control. How this pattern of default-executive coupling unfolds across the lifespan and relates to cognitive performance remain unclear, and, importantly, longitudinal data are scarce. Here, we tested the main hypotheses of this DECHA model across the lifespan (ages 6-81) using measures of longitudinal changes in cognitive control function over 3.5 years and functional connectivity (FC) during incidental encoding and retrieval of episodic memories.

While DECHA does not directly address development, it is broadly known and agreed that cognitive control increases with maturation (Best & Miller, 2010; Constantinidis & Luna, 2019; Luna et al., 2015) and decreases with aging (Ferguson et al., 2021; Manard et al., 2014; Xia et al., 2022). Thus, it is of interest to investigate hypotheses of aging in relation to development (Walhovd et al., 2020). Specifically, DECHA consists of two central hypotheses. The first hypothesis states that general cognitive decline in aging is accompanied by increased functional coupling between the default mode network (DMN) and the left dorsolateral prefrontal cortex (dlPFC). The DMN is a large-scale brain network which includes medial temporal lobe regions, involved in episodic memory, mental simulation and future imagination (Andrews-Hanna et al., 2010; Daselaar et al., 2009; Kim, 2010; Kizilirmak et al., 2023). The dlPFC is involved in prioritizing task-relevant information and likely plays a crucial role in cognitive control (Jiang et al., 2018; Petrides, 2005; Turnbull et al., 2019).

In support of this first hypothesis of DECHA, several cross-sectional studies have shown a pattern of greater dlPFC-DMN coupling in older relative to younger adults during tasks of cognitive control (Dixon et al., 2018; Grady et al., 2016; A. C. Murphy et al., 2020). A foundational study demonstrated that higher FC between the left dlPFC and key nodes of the DMN was associated with worsened goal-oriented planning behavior in adults between 63 and 78 years of age, while the same pattern of FC was not found in younger adults (18-27 years), indicating different levels of cognitive control relating differentially to this pattern of functional connectivity (Turner & Spreng, 2015). However, in the context of episodic memory, only one study has associated FC between the DMN and a network including the dlPFC, and episodic memory performance in adults over 70 years of age (Zhukovsky et al., 2023). This association was derived from resting-state fMRI data within a limited age range and in a clinical sample. Hence, it remains unclear how dlPFC-DMN coupling during episodic memory operations evolve with age in normal aging.

The second main hypothesis of DECHA is that the age-related increase in dlPFC– DMN coupling co-occurs with a decline in cognitive control resources (Spreng & Turner, 2019), a hallmark of cognitive aging (Park & Reuter-Lorenz, 2009). Cognitive control refers to the set of processes necessary for planning and executing goal-directed behaviors and for inhibiting irrelevant or competing responses, which in turn likely impact episodic memory performance (Anderson et al., 2016; Badre & Wagner, 2007; Kim, 2020; Ray et al., 2019). According to DECHA, diminished cognitive control and increased default–executive coupling jointly contributes to broader cognitive decline. This loss of control resources is also thought to be accompanied by a shift toward greater reliance on existing knowledge structures, a process referred to as the *semanticization* of cognition (e.g., Park et al., 2002; Singh-Manoux et al., 2012). Lifespan studies of cognitive control have shown age-related differences cross-sectionally (Ferguson et al., 2021) and longitudinally (Adólfsdóttir et al., 2017), including in a partially overlapping sample (Fjell et al., 2017). However, previous studies have not investigated the link between dlPFC-DMN coupling and longitudinal changes in cognitive control, which is critical in determining whether age-related differences in FC associate with a decline in cognitive control resources.

Cognitive control shows consistent improvement from childhood into early adulthood (Best & Miller, 2010; Constantinidis & Luna, 2019; Luna et al., 2015) and some findings suggest that this may happen in parallel with dlPFC-DMN decoupling. For instance, a longitudinal study of participants between the ages of 10 and 13 found decreased resting-state FC between the DMN and networks of cognitive control (including the left dlPFC) (Sherman et al., 2014), consistent with increasing functional specialization of brain networks during development (Grayson & Fair, 2017). However, other studies with broader age ranges have reported no association between age and dlPFC-DMN connectivity (Betzel et al., 2014; Breukelaar et al., 2020). These inconcistencies leave open the question of how default-executive coupling evolves across the lifespan.

Although much of the existing research on DECHA has focused on cognitive control (Dixon et al., 2018; Grady et al., 2016; A. C. Murphy et al., 2020; Turner & Spreng, 2015), the model is intended to explain broader patterns of cognitive decline in aging through underlying brain network dynamics. Episodic memory is a domain that undergoes marked changes across the lifespan and may also be influenced by default-executive network dynamics. Memory performance improves through childhood and adolescence (Selmeczy & Ghetti, 2025; Shing & Lindenberger, 2011), paralleling structural maturation of the medial temporal lobe and dlPFC (Gogtay et al., 2004; Sowell et al., 2003). Decline in episodic memory have been observed as early as the third decade of life in both cross-sectional and longitudinal studies (Capogna et al., 2022; Salthouse, 2009; Vidal-Piñeiro et al., 2019), although considerable inter-individual variability exists and other studies suggest later onset of decline (Nyberg et al., 2012). While the brain mechanisms underlying this variability are likely complex and multifaceted (Nyberg, 2017; Small, 2001; Tromp et al., 2015), DECHA offers a specific and testable mechanism to help explain these variations in memory performance using a task-based functional imaging approach (Mooraj et al., 2025).

In the present study, we used fMRI during episodic memory encoding and retrieval to test the dlPFC-DMN FC with age in a lifespan sample, and whether this is associated with changes in cognitive control up to 10 years earlier. We also explored the specificity of dlPFC-DMN FC by comparing the dlPFC-DMN coupling to dlPFC coupling with 16 other functional networks (Schaefer et al., 2018). We operationalized cognitive control using completion time in the D-KEFS Stroop inhibition/switching task (Delis et al., 2012), typically associated with cognitive control abilities (Spreng and Turner 2019). This Stroop subtest captures age-related longitudinal decline in cognitive control well (Clark et al., 2012; Fjell et al., 2017), and is likely dependent on the dlPFC during interference control (Banich, 2019; Liu et al., 2006; Milham et al., 2003). To ease the interpretation of likely non-linear lifespan trajectories, we divided the sample into a youth (6-29 years) and an aging subsample (30-81 years) based on literature showing episodic memory decline after the age of 30 (Capogna et al., 2022; Salthouse, 2009; Vidal-Piñeiro et al., 2019). We hypothesized that (i) dlPFC-DMN FC would be greater with age in the aging subsample and inversely, lesser with age in the youth subsample. (ii) We further hypothesized that longitudinal increase in Stroop completion time would be associated with higher levels of dlPFC-DMN FC. (iii) Finally, we expected that such higher levels of dlPFC-DMN FC would be associated with lower memory performance.

## 2 Methods

### 2.1 Participants

Written informed consent was obtained from all adult participants and from parents or other legal guardians for participants below the age of 18. The study was approved by the Regional Ethical Committee of South Norway and conducted in accordance with the Helsinki declaration. A total of 552 healthy participants (371 females, 181 males, mean age = 39.9 years, SD = 18.5, age range = 6–81) were part of the cross-sectional fMRI sample (**Table 1**). Participants who successfully performed the fMRI task underwent screening to ensure they had no history of neurological disorders or chronic illness. They needed to have right-hand dominance and not be taking any medication that may impact the functioning of their nervous system. Additional exclusion of participants was based on neuropsychological criteria including: MMSE score for participants over 50 < 26 (Folstein et al., 1975), IQ score < 85 (Wechsler, 1999), and Beck’s Depression Inventory (BDI) > 20 (Beck et al., 1988). A total of 452 out of the 552 participants in the cross-sectional sample had at least one neuropsychological assessment, including Stroop, at the same time as fMRI collection. A total of 198 of these had in addition completed Stroop one to six times before fMRI collection (**Figure 1B**). The mean (SD) interval back in time from fMRI acquisition was 3.5 (2.66) years, max interval 10 years (**Table 2**). See **Figure 1A** for a schematic of the study design.

**Figure 1.**
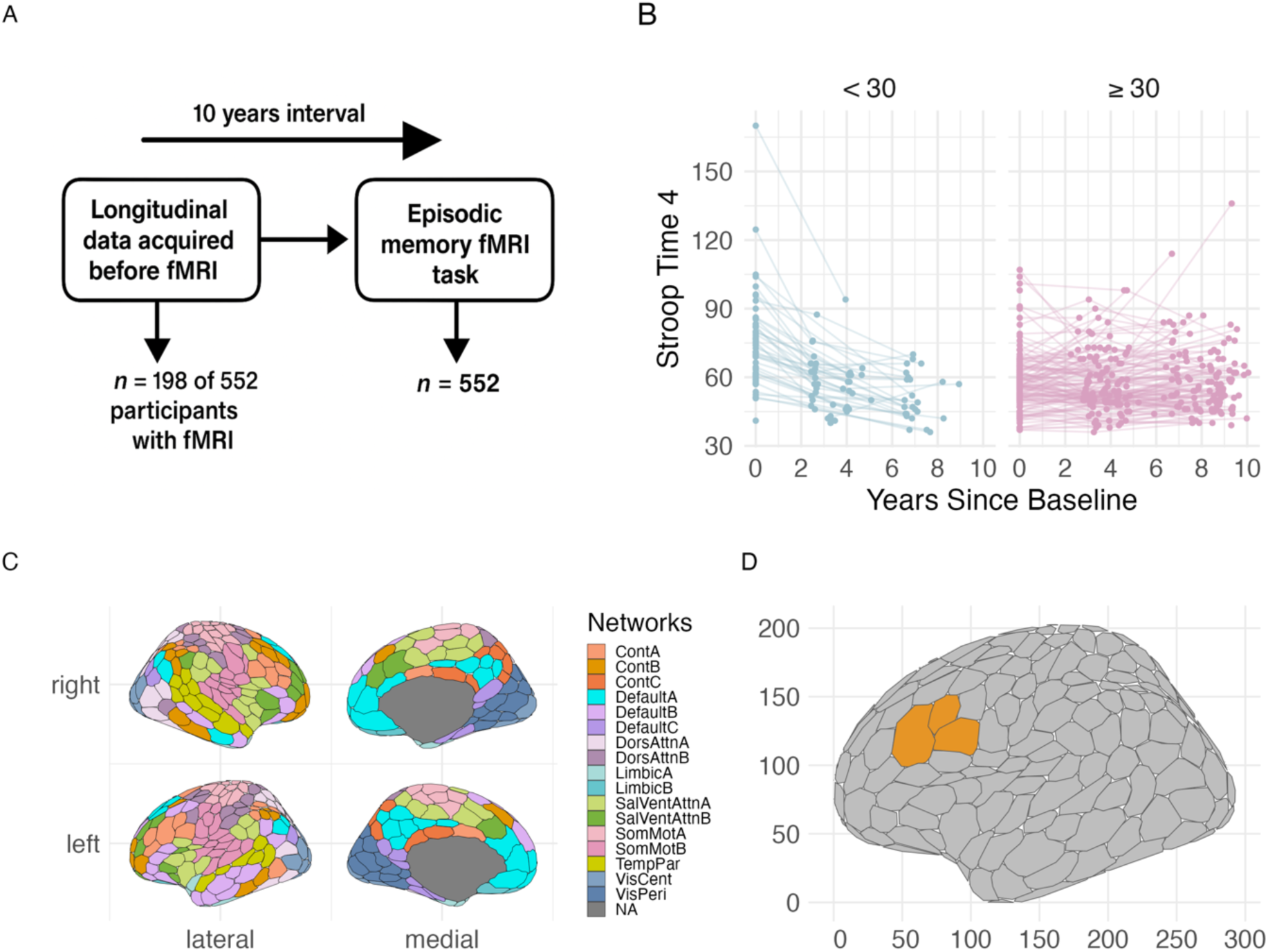
A) Schematic of study timeline. B) Spaghetti plot over participants with multiple Stroop measurements before fMRI acquisition, up to 10 years interval, divided into ages </> 30 years. Points display measurement and lines indicate time between measurements. The last point for each participant is the closest Stroop measurement before fMRI acquisition. C) Schaefers 17 networks 400 parcellations atlas used to create functional connectivity estimations. D) Seed regions of left dlPFC found in Control A network from Schaefers 17 networks atlas.

**Table 1.**

| Variables | Mean (SD) | Range | N obs |
| --- | --- | --- | --- |
| **Age Group < 30** |  |  |  |
| Age (MRI) | 22.39 (6.23) | 6 – 29 | 238 |
| IQ | 111.94 (9.47) | 89 – 137 | 154 |
| MMS | 29.21 (0.96) | 26 – 30 | 150 |
| BDI | 4.74 (3.7) | 0 – 14 | 31 |
| Sex (females/males) |  |  | 159/79 |
| **Age Group ≥ 30** |  |  |  |
| Age (MRI) | 51.30 (14.76) | 30 – 81 | 314 |
| IQ | 117.19 (10.19) | 90 – 138 | 212 |
| MMS | 29.03 (0.98) | 26 – 30 | 279 |
| BDI | 4.23 (3.7) | 0 – 16 | 41 |
| Sex (females/males) |  |  | 212/102 |

Cross-sectional fMRI sample demographics.

**Table 2.**

|  | **Age Group < 30** |  |  |  |  |  |
| --- | --- | --- | --- | --- | --- | --- |
| Visit Number | 1 (N = 22) | 2 (N = 17) | 3 (N = 27) | 4 (N = 0) | 5 (N = 0) | 6 (N = 0) |
| Age Mean (SD) | 12.57 (3.18) | 17.06 (4.41) | 21.88 (5.80) |  |  |  |
| Age Range | 8.89 – 24.60 | 11.55 – 26.62 | 15.67 – 29.83 |  |  |  |
| Stroop Mean (SD) | 74.84 (20.14) | 57.57 (26.62) | 51.19 (9.92) |  |  |  |
| Stroop Range | 41 – 170 | 36 – 94 | 37 – 70 |  |  |  |
| Female | 27 (45.8) | 20 (45.5) | 15 (55.6) |  |  |  |
| Males | 32 (54.2) | 24 (54.5) | 12 (44.4) |  |  |  |

**Age Group $\geq 30$**
| Visit Number | 1 (N = 1) | 2 (N = 12) | 3 (N = 106) | 4 (N = 2) | 5 (N = 5) | 6 (N = 6) |
| --- | --- | --- | --- | --- | --- | --- |
| Age Mean (SD) | 56.03 (11.61) | 56.54 (11.09) | 61.93 (10.96) | 78.78 (7.60) | 76.25 (2.70) | 75.03 (2.98) |
| Age Range | 30.83 – 78.59 | 30.93 – 77.27 | 35.34 – 80.89 | 70.40 – 81.15 | 71.76 – 78.67 | 72.10 – 79.97 |
| Stroop Mean (SD) | 56.64 (13.57) | 56.90 (12.82) | 59.91 (15.18) | 61.50 (14.85) | 61.20 (16.53) | 68.83 (10.17) |
| Stroop Range | 37 – 107 | 36 – 98 | 39 – 136 | 51 – 72 | 44 – 86 | 52 – 80 |
| Female N (%) | 68 (56.7) | 66 (57.4) | 64 (59.8) | 1 (50) | 4 (20) | 5 (83) |
| Males N (%) | 52 (43.3) | 49 (42.6) | 43 (40.2) | 1 (50) | 1 (20) | 1 (16.7) |

Longitudinal sample demographics of participants completing neuropsychological testing up to 10 years before fMRI data collection. Visit number indicates how many participants had up to $n$ neuropsychological measurements. Stroop refers to Stroop completion time in seconds.

### 2.2 Out-of-scanner cognitive measures

For the Stroop color-word interference task, a modified version of the Stroop task was administered (Delis et al., 2001). Out of the four conditions in the task, (1) color naming, (2) reading, (3) inhibition and (4) inhibition/switching, we used the inhibition/switching condition for our analyses. In this condition, the participants must alternate between reading the colored word and naming the ink color of a word. Specifically, this task measures the response time of naming the color of a word that is either congruent or incongruent with the words’ semantic content and, thereby, the capacity to override the automatic response of reading a word. This design introduces additional demand on cognitive control resources and is shown elsewhere to relate to longitudinal changes (Fjell et al., 2017). The participant is required to complete 48 trials in the condition as fast as possible, where correct and incorrect responses are also noted. To examine age-related changes in semantic knowledge, the Wechsler Abbreviated Scale of Intelligence (WASI) (Wechsler, 1999) vocabulary task was administered. In this task the participant is read a list of words with increasingly difficulty by an examinator, to which they must define the word. We used the raw vocabulary scores to examine age trajectories. Word difficulty is calibrated by age in development (i.e., participants 6-8 years of age starts with an easier word than participants 12 years of age).

### 2.3 Experimental design and procedure

The fMRI experiment included an incidental encoding task and a retrieval phase after approximately 90 min, both in the scanner. Details of the experimental design have been described by others (Sneve et al., 2015; Vidal-Piñeiro et al., 2019). Briefly, during the encoding phase participants were presented with 300 “everyday objects” in black and white drawings. The questions “Can you eat it?” or “Can you lift it” followed the stimulus presentation. Participants were instructed to respond “yes” or “no” with a button press. The assignment of button presses was counterbalanced across participants. During the retrieval phase new and old items were presented, preceded by the questions “Have you seen this item before?”. If no (new item), another item was presented. If yes (old item) the following question was presented: “Can you remember what to do with the item?”. Again, if no, the trial ended. If the participant answered yes, the following question was presented: “Were you supposed to eat it or lift it?”. A forced choice associated with the two actions during encoding (eat/lift) was presented and answered with a button press. Classification of responses to old items was: (1) source memory (“Yes” response to Question 1 and Question 2, and correct answer to Question 3), (2) item memory (“Yes” response to Question 1 and either “No” to Question 2 or incorrect answer to Question 3), (3) miss (incorrect answer to Question 1). New items were classified either as (4) correct rejections or (5) false alarms. We used (1) source memory as the measure of episodic memory in our analyses.

### 2.4 MRI acquisition

Data was acquired using a 3T MRI (Siemens Skyra Scanner, Siemens Medical Solutions, Germany) and a 20-channel Siemens head-neck coil at Rikshospitalet, Oslo University Hospital. Functional imaging parameters remained consistent throughout all fMRI runs, with a BOLD-sensitive T2*-weighted EPI sequence measuring 43 transversally oriented slices with no gap (TR = 2390 ms, TE = 30 ms, f lip angle = 90◦, voxel size = 3 × 3 × 3mm3, FOV = 224 × 224 mm2, interleaved acquisition; generalized autocalibrating partially parallel acquisitions acceleration factor [GRAPPA] = 2). Each encoding run generated 134 volumes, and 3 dummy volumes were acquired at the start of each fMRI run to avoid T1 saturation effects. Anatomical T1-weighted (T1w) magnetization-prepared rapid gradient echo (MPRAGE) images were obtained using a turbo field echo pulse sequence consisting of 176 sagittally oriented slices, while a standard double-echo gradient-echo field map sequence was acquired for distortion correction of the echo planar images. Visual stimuli were presented on a 32-inch LCD monitor, and participants responded using the ResponseGrip device (NordicNeurolab, Norway). Auditory stimuli were presented through the scanner intercom and delivered to the participants’ headphones.

### 2.5 fMRI preprocessing

Data were preprocessed using fMRIPrep (v. 1.5.3; Esteban et al., 2019, 2020) and organized according to the Brain Imaging Dataset Specification (BIDS). The T1- weighted (T1w) anatomical image was corrected for intensity nonuniformity (INU) and used as the reference throughout the pipeline. The T1w-reference was skull-stripped, and all functional runs were co-registered to this reference image using boundary-based registration with six degrees of freedom.

Susceptibility-induced distortion correction was performed using a custom implementation of the TOPUP technique (https://github.com/markushs/sdcflows/tree/topup_mod; Andersson et al., 2003). Each BOLD run underwent slice-timing correction and motion correction, with motion parameters estimated from rigid body transformations. Denoising was performed using ICA-AROMA (Pruim et al., 2015), incorporating mean white matter and cerebrospinal fluid time series as nuisance regressors, along with the six motion parameters. Detrending and high-pass filtering (0.008 Hz) were applied using a fifth-order Butterworth filter, orthogonalized to the nuisance regressorss (Hallquist et al., 2013).

Global signal regression (GSR) was used to reduce global non-neuronal signals which can spuriously inflate functional connectivity estimates (Li et al., 2019; Power et al., 2014). Although GSR remains controversial due to its potential to introduce artificial anticorrelations and to some degree remove meaningful neural signals (K. Murphy et al., 2009; Saad et al., 2012), it has also been shown to enhance the specificity of functional networks focused on regionally specific connectivity (Fox et al., 2009; Li et al., 2019). In line with recent recommendations by Bijsterbosch et al. (2020), all analyses were also conducted without GSR.

### 2.6 Functional connectivity estimation

Functional connectivity was estimated with correlational psychophysiological interaction (cPPI) for each individual native space, using a ROI-based approach using partial correlations, controlling for task-unrelated activity (Fornito et al., 2012). Because of the way cPPI is estimated, it includes both task-related activity and intrinsic background activity resembling resting-state FC. As such, FC values derived from cPPI across tasks (encoding, retrieval) are likely to be highly similar and closely resemble steady-state FC rather than traditional PPI measures. The cerebral cortex was divided into 400 nodes divided into 17 functional connectivity networks based on the Schaefer et al. parcellation (Schaefer et al., 2018). In addition, three dlPFC nodes from the ControlA network in the left hemisphere were extracted and were treated as their own network. The cPPI procedure for the same sample during encoding has previously been described (Capogna et al., 2022; Grydeland et al., 2025). Briefly, boxcar functions were used to construct the psychological time-series, reflecting 2 s events from the start until the end of each trial. Next, deconvolving BOLD time-series from each region the PPI terms were created (Gitelman et al., 2003), and multiplied point-by-point with the psychological time-series, followed by re-transforming the PPI terms to BOLD via convolution with a canonical 2-gamma HRF. Finally, cPPI matrices were constructed through partial Pearson’s correlation analysis and the PPI terms are adjusted for the regions’ BOLD timeseries and the HRF-convolved psychological timeseries. As described elsewhere (Raud et al., 2023), cPPI estimation of the retrieval phase used all the time-points of all three questions to calculate the regressors. However, only the estimates for question 1 were utilized in forming the cPPI matrices. This approach ensures the inclusion of all trials only once, while capturing variability from all task events. The correlation coefficients obtained were Fisher-transformed into z-values. Prior to higher-order analyses, average within- and between-network connectivity was computed for the 17 networks and the three dlPFC nodes. This was done by averaging the Fisher z-transformed connectivity values between all pairs of ROIs within and across networks, based on the cPPI matrices.

### 2.7 Statistical analysis

#### Lifespan and Longitudinal Cognitive Trajectories

To examine age-related patterns in cognitive performance, we used generalized additive models (GAMs) for cross-sectional data and generalized additive mixed models (GAMMs) for longitudinal data, which allow flexible modeling of non-linear and linear trends while accounting for fixed and random effects (Sørensen et al., 2021). Lifespan trajectories of in-scanner episodic memory were modeled using a GAM, while trajectories of cognitive control (Stroop inhibition/switching completion time) and vocabulary performance were modeled using GAMMs.

Longitudinal change in cognitive control and vocabulary was assessed using separate linear mixed effects models for the youth and aging subsamples. For cognitive control, models included baseline age, sex, Stroop reading and color naming scores, a test-retest indicator (0 vs. ≥ 1 previous test), and an interaction between baseline age and time between assessments. Time was modeled as a random slope for each participant. Vocabulary models followed the same structure, excluding Stroop-specific covariates.

#### Cross-sectional FC–Age Associations

To test age-related variation in dlPFC–DMN FC, we used linear regression models with FC as the dependent variable separately for the youth and aging subsamples. Predictors included age, sex, in-scanner movement, and subject-wise average cortical FC. We focused on the DMNa subsystem from the Schaefer et al. (2018) atlas due to its relevance for distractibility and its inclusion of medial prefrontal cortex (mPFC) and posterior cingulate cortex (PCC) nodes (Andrews-Hanna et al., 2010; Turner & Spreng, 2015).

#### Longitudinal FC–Cognitive Control Associations

To assess whether dlPFC–DMNa FC predicted change in cognitive control, we used GAMMs with Stroop inhibition/switching completion time as the outcome. Models included the interaction between FC and time, along with sex, in-scanner movement, Stroop reading and color naming scores, and subject-wise average cortical FC, following a level-change modeling approach (Fjell et al., 2023). Analyses were conducted separately for the youth and aging subsamples across all dlPFC–network connections.

#### Cross-sectional FC–Memory Associations

We used linear regression models to examine associations between FC and memory performance, including interactions with age. All models included age, sex, in-scanner movement, and subject-wise average cortical FC. Interaction terms were tested in separate models.

#### Multiple Comparison Correction

To assess network specificity, all relevant terms (main and interaction) from the models testing dlPFC connectivity with each of the 16 other networks were corrected for multiple comparisons using a 5% false discovery rate (FDR; Benjamini & Hochberg, 1995). All uncorrected main FC findings can be found in Supplementary Material (SM), section 3.

## 3 Results

### 3.1 Age associations for memory, cognitive control, and semantic knowledge

**Figure 2A** shows the age-trajectory of the full lifespan sample of episodic memory performance (*F* = 29.19, edf = 6.73, p < 0.001). The age-memory trajectory showed that performance in the task stabilizes between 20 and 30 years of age, a decline is noticeable from around the 30-year mark, followed by even larger differences starting in the fifth decade of life.

**Figure 2.**
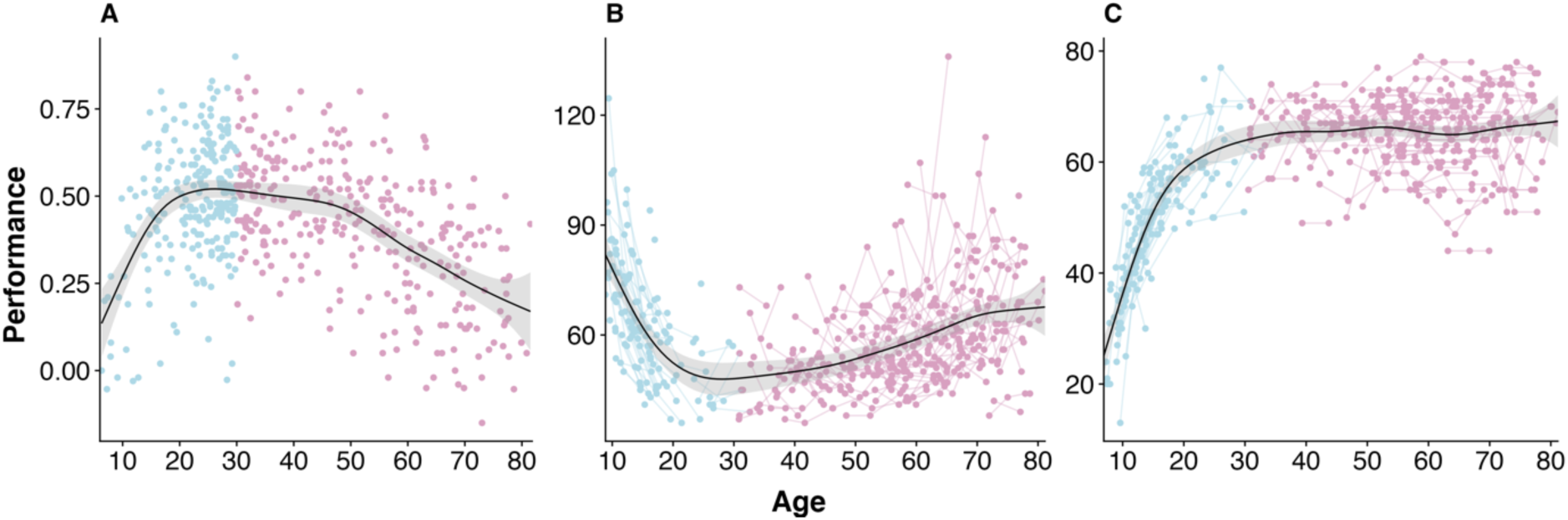
Lifespan trajectories of cognitive measures. A) Cross-sectional, in-scanner source memory performance, B) Longitudinal, out-of-scanner cognitive control (Stroop inhibition/switching completion time) and C) vocabulary. Blue color indicates age under 30 and purple color indicates the ages over 30. Lines between points indicate multiple measurements over time for a participant.

A central hypothesis of DECHA is that the age-related increase in dlPFC-DMNa FC is accompanied by an age-related decline in cognitive control, and a suggested shift towards a greater reliance on prior knowledge, including semantic abilities. To investigate this predicted shift in cognition, we used longitudinal measures of cognitive control and semantic abilities taken at the same time as the fMRI and up to 10 years earlier (mean interval = 3.5 years, mean number of measurements per participant = 2.66). Across the full age-range, there was a non-linear relationship between age and completion time during the Stroop inhibition condition (**Figure 2B**, *F* = 11.54, edf = 5.255, *P* < 0.001). A non-linear relationship with age was also observed for vocabulary (**Figure 2C**, *F* =109.1, edf = 8.307, *P* < 0.001).

To aid the interpretation of the non-linear lifespan associations, we divided the sample in 2 subsamples, a youth group and an aging group, based on evidence of episodic memory decline after 30 years (**Figure 2A**, cutoff = 30 years). For the aging subsample, analyses revealed a positive interaction between age and change, with Stroop completion time change being larger with higher age (**SM** **Figure 1A**, b = 1.62, t = 2.83, p = 0.005). Vocabulary did not show an interaction between age and change (**SM** **Figure 1B**, b = -0.08, t = -0.33, p = 0.7).

As a control analysis (**SM** **Figure 2)**, we tested whether changes in Stroop completion time associated with memory performance for the aging subsample. Using GAMM, we found that an increase in Stroop completion time over time was associated with poorer memory performance (edf = 1.00, F. = 6.12 p = 0.01).

### 3.2 Memory Functional Connectivity and Age

#### 3.2.1 dlPFC-DMNa FC and Age

We then examined the main hypothesis of DECHA, namely higher FC between the dlPFC and DMNa with higher age in the aging group, as well as our extension to the youth group, where we expected the inverse relationship. **Figure 3A** and **3B** show lifespan trajectories during encoding and retrieval, respectively (**SM** **Figure 3** shows linear age relationships for dlPFC-DMNa FC per age-group). In the aging group, linear regression models showed that higher age was associated with higher FC during both encoding (b = 0.20, t = 3.2, p = 0.001) and retrieval (b = 0.23, t = 3.66 p < 0.001). In the youth group, higher age was associated with lower FC during retrieval, (b = -0.14, t = -1.98, p = 0.04), but not during encoding (b = -0.07, t = -1.09, p = 0.2). This main result was highly similar in both groups when repeating the analyses using FC without performing GSR (**SM** **Figure 6).**

**Figure 3.**
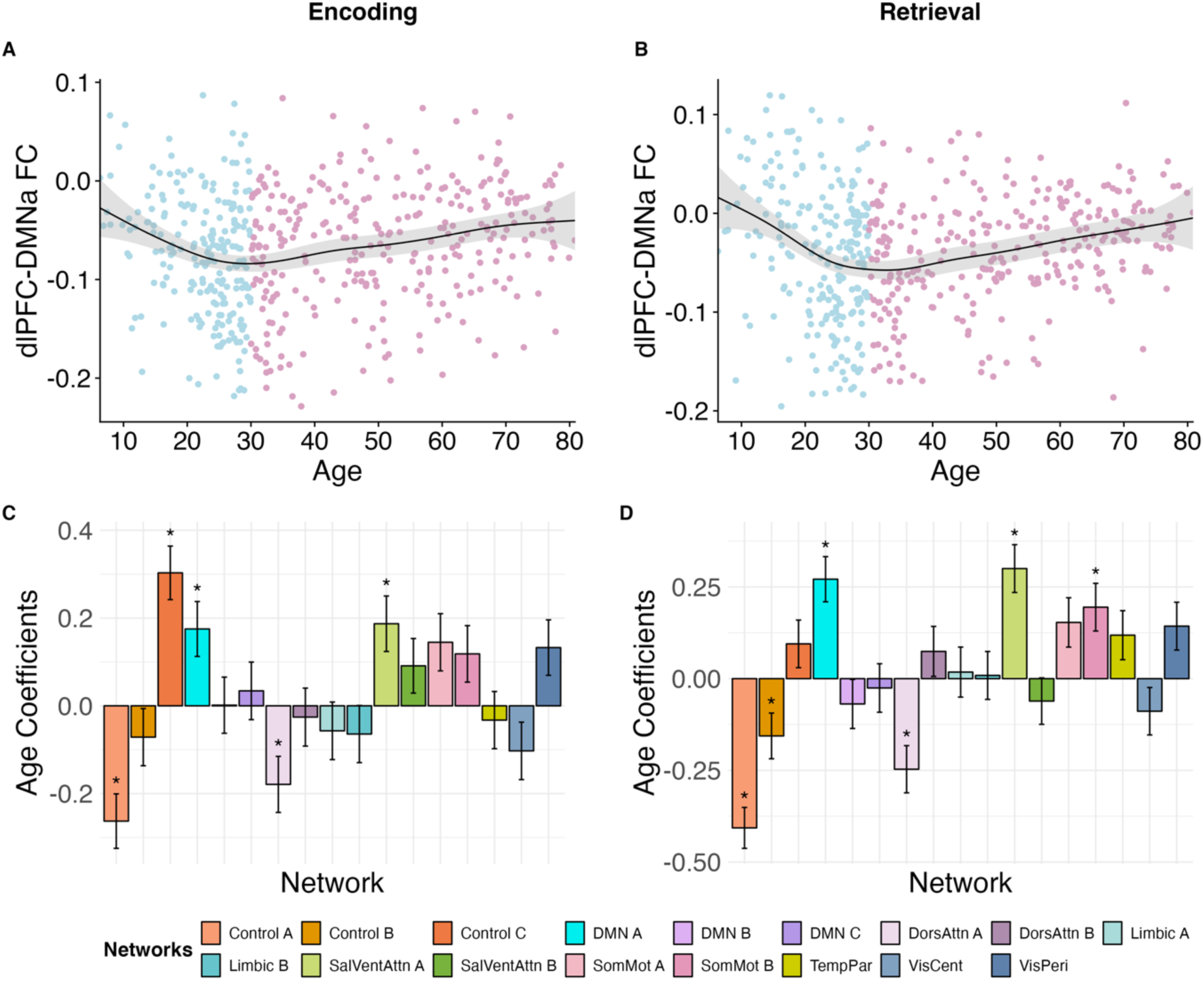
Cross-sectional relationships between age and FC between the dlPFC and DMNa. movement. **A)** Lifespan trajectory for dlPFC-DMNa FC during encoding and **B)** retrieval. Blue dots represent the youth subgroup, and the purple dots represent the aging subgroup. **C)** All significant age associations for the aging subsample during encoding, and **D**) during retrieval.

#### 3.2.2 Specificity of dlPFC-DMNa FC

To test whether the association between dlPFC and DMNa with age was a specific feature of dlPFC connections, we tested age-related FC between the dlPFC and the remaining 16 networks. As shown in **Figure 3C**, during encoding for the aging subsample, age was related to higher FC between dlPFC and ControlC (b = 0.30, t = 4.97, FDRp < 0.001), SalventAttnA (b =0.18, t = 2.95, FDRp = 0.03), and lower FC between dlPFC and ControlA (b = -0.26, t = -4.23, FDRp = 0.001), and DorsAttnA (b = -0.17, t = -2.80, FDRp = 0.01). At retrieval (**Figure 3D**), positive relationships with age were observed for SalventAttnA (b = 0.29, t = 4.60, FDRp = 0.001), as well as SomMotB (b = 0.19, t = 2.99, FDRp = 0.01). Negative relationships between FC and age were also observed, between dlPFC and ControlA (b = -0.40, t = -7.29, FDRp < 0.001), DorsAttnA (b = -0.24, t = -3.84, FDRp = 0.002) and ControlB (b = -0.15, t = - 2.50, FDRp. = 0.03), respectively. Analyses using FC without GSR demonstrated highly similar age relationships for the networks presented here (**SM** **Figure 7**, ControlC, SalventAttnA, SomMotA-B, and ControlA [retrieval]). However, there were some exceptions: ControlA and DorsAttnA at encoding, as well as ControlB at retrieval, were not significant, and the DorsAttnA network at retrieval showed an age effect in the opposite direction (positive with GSR). Age effects was also seen for SalventAttnB, TempPar, VisCent, VisPeri at encoding, and DorsAttnB, SomMotA, TempPar, VisCent, VisPeri at retrieval.

For the youth subgroup, no dlPFC-network FC associations with age survived correction for multiple comparisons. Although dlPFC-DMNc FC at encoding did not survive correction for multiple comparisons with GSR, the direction of the slope was consistent with and without GSR. Without GSR, the FC at encoding between dlPFC and DMNc survived FDR correction (b = -0.12, t = -3.09, FDRp = 0.03).

### 3.3 Associations Between Memory FC and Preceding Change in Cognitive Control

Using GAMM, we examined whether change in cognitive control related to the level of episodic memory-related FC. As shown in **Figure 4** (upper left panel) in the aging group, higher FC between left dlPFC and DMNa related to an increase in Stroop completion time during encoding (edf = 3.26, F = 2.11 p = 0.04), but not during retrieval (edf = 1.00, F = 0.14 p = 0.76). This result was replicated in the FC analysis without GSR (**SM** **Figure 8**).

**Figure 4.**
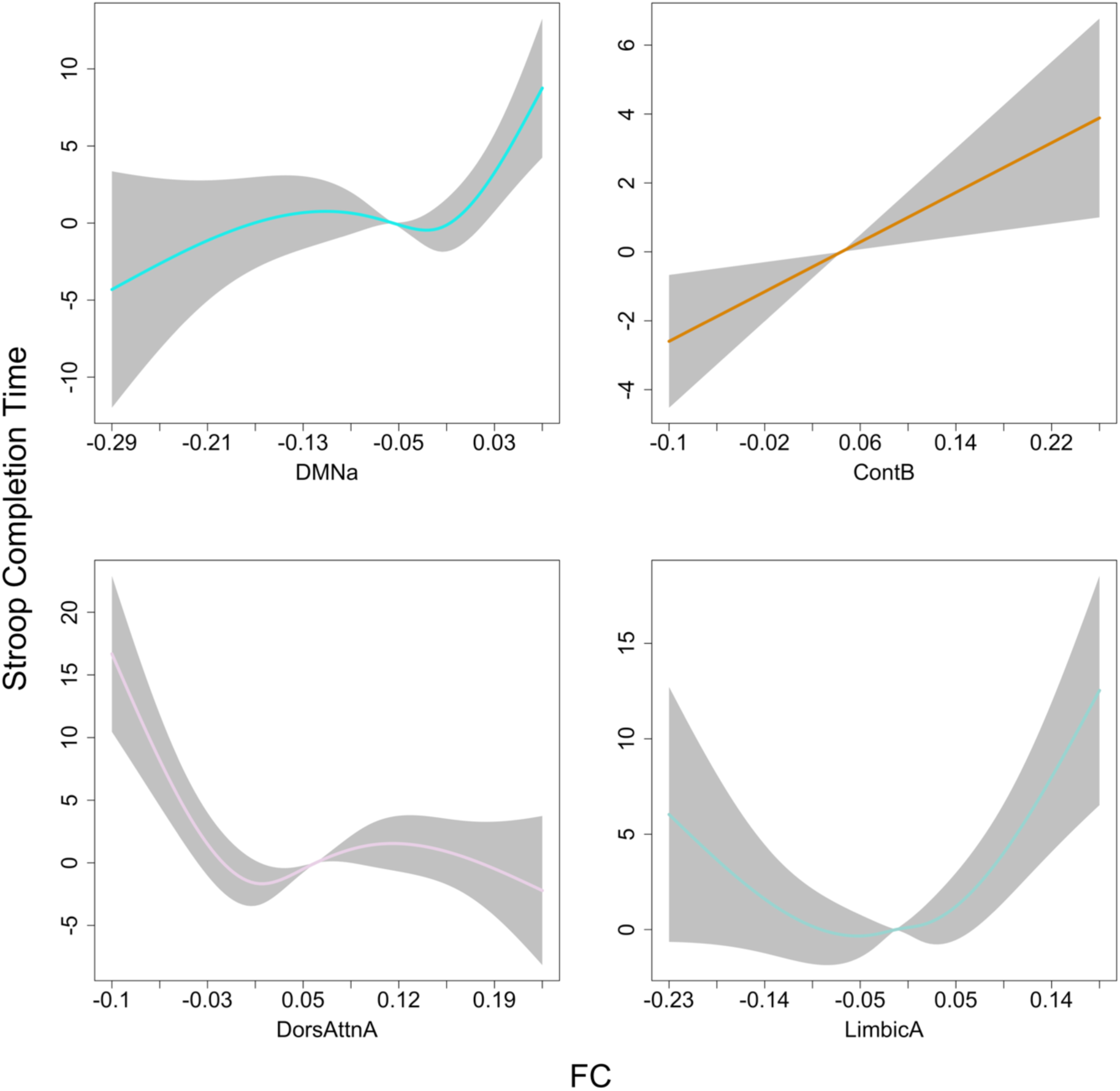
Longitudinal change in cognitive control, as measured by Stroop completion time, varying with dlPFC-to-network FC during encoding. On the y-axis, change in completion time is represented, with positive values indicating an increase in completion time. The x-axis reflects the level of functional connectivity. **Upper left panel =** DMNa FC during encoding, **upper right panel =** ControlB FC during encoding, **lower left panel =** DorsAttna during encoding, **lower right panel =** LimbicA FC during encoding for the aging subsample.

To test the specificity of dlPFC-DMNa FC relationship with cognitive control change, we tested how dlPFC coupling with other networks associated with cognitive control change. During encoding, increased Stroop completion time correlated with higher FC between left dlPFC and the ControlB network (**Figure 4**, upper right panel, edf = 1.00, F = 7.78 p < 0.01, FDRp = 0.04), the LimbicA displayed an U-shape (**Figure 4**, lower right panel, edf = 3.17, F = 5.10, p = 0.003, FDRp = 0.01), while decreased Stroop inhibition completion time correlated with higher FC between dlPFC and the DorsAttnA network (**Figure 4**, lower left panel, edf = 3.57, F = 8.91 p < 0.001, FDRp = 0.008). Only dlPFC-DorsAttnB FC was significantly associated with longitudinal change in cognitive control during retrieval without GSR (**SM** **Figure 8**). Except for dlPFC-DMNa, no other association was found during encoding after multiple comparison corrections.

Similar analysis in the youth subsample showed a weak, non-significant association for DMNa during retrieval (edf = 2.00, F = 2.35, p = 0.09), but not during encoding (edf =1.00, F = 0.57 p = 0.44).

### 3.4 Relationships with Memory Performance

Finally, we assessed how task memory performance linked with dlPFC-network FC. In the aging subgroup (**SM** **Figure 4)**, memory performance did not associate with dlPFC-DMNa FC at encoding (b = 0.02, t = 0.42, p= 0.66) or at retrieval (b = 0.08, t = -1.55, p.= 0.12). For the other dlPFC couplings, FC between dlPFC and ControlA at retrieval was positively associated with better memory performance after correction for multiple comparisons (b = 0.03, t = 3.16, FDRp = 0.02). For the youth subgroup, we did not observe any associations between FC and memory performance. Similar non-significant results were observed when using FC without GSR.

## 4 Discussion

In line with the DECHA framework, we found that FC between DMN and dlPFC during memory operations showed an age-related difference from young adulthood into older age. We also extended this framework by showing inverse age relationships in a youth group aged 6-29 years. Further, in adults aged 30-81 years, greater dlPFC-DMN FC was associated with longitudinal decline in cognitive control, but not with memory performance, which instead was found between dlPFC and other control network regions. Similarly, age-related FC patterns were also observed between the dlPFC and other networks, particularly salient attention and control subnetworks. Taken together, our results support the main hypotheses of DECHA, while also suggesting that these hypotheses at least partly fit a large lifespan framework where also broader dlPFC connectivity profiles may play prominent roles as neural mechanisms behind cognitive decline in cognitive control and episodic memory.

### 4.1 How does functional connectivity between the default-mode network and the dlPFC vary across the lifespan?

In line with previous results in adults, greater age was related to higher FC between left dlPFC and a DMN subnetwork (Grady et al., 2016; Turner & Spreng, 2015; Zhukovsky et al., 2022). This DMN subnetwork is one of three DMN subnetworks (Yeo et al., 2011) and includes the DMN regions highlighted as important in DECHA, including the medial prefrontal cortex, posterior cingulate cortex, and the angular gyrus (Turner & Spreng, 2015). Our results expand the existing literature by demonstrating a U-shape of dlPFC-DMNa FC across the lifespan. We observed higher levels of FC with higher age in the aging group and lower FC with age in the youth subgroup, specifically during retrieval. DECHA suggests that increased FC between the DMN and dlPFC accompanies a cognitive mode characterized by more mind-wandering and distractibility (Spreng & Turner, 2019). This shift is attributed to the putative involvement of the dlPFC in prioritizing neural processing (Petrides, 2005; Jiang et al., 2018; Turnbull et al., 2019) and the DMN in mind-wandering and self-referential associations (Andrews-Hanna et al., 2010, 2014). Thus, according to DECHA, the coupling of these networks predicts a higher degree of distractibility and stronger reliance semanticized knowledge (Spreng & Turner, 2019). This idea of distractibility varying with dlPFC-DMNa FC is in line with our other finding of reduced cognitive control (measured as an increase in Stroop completion time) relating to higher FC between dlPFC and the DMN subnetwork during encoding.

However, as several other dlPFC-network connections also displayed age relationships, the age-related differences in coupling between the dlPFC and DMN does not seem to be DMN specific. This finding supports the growing body of evidence where age-related functional recoupling is characterized by more between- FC from prefrontal areas to other regions in the brain with age (Andrews-Hanna et al., 2007; Damoiseaux et al., 2008; Kizilirmak et al., 2023), also demonstrated in a similar sample and the same fMRI-task as the one used in the present study (Capogna et al., 2022).

We found a significant negative age-related slope in the dlPFC-DMN connectivity during retrieval in the youth subsample, whereas encoding did not show a significant age association, but still exhibited a negative age relationship. In the literature, the emerging evidence of dlPFC-DMN coupling in development remain inconclusive.

Declining resting-state FC between networks of cognitive control (including the left dlPFC) and the DMN has previously been demonstrated in a longitudinal study between the ages of 10 and 13 (Sherman et al., 2014), while studies of resting-state FC in longitudinal samples between the ages 8 and 39 (Breukelaar et al., 2020) and a large cross-sectional lifespan sample (Betzel et al., 2014) did not find correlation between DMN and cognitive control networks FC with age. Our findings indicates that dlPFC-DMN FC evolve throughout the stages of childhood, adolescence, and young adulthood.

### 4.2 How does longitudinal changes in cognitive control vary with level of FC?

In aging, we found that dlPFC-DMN FC was positively related to reductions in cognitive control as measured by the Stroop inhibition/switching completion time. In line with DECHA, this result supports the proposition that loss of cognitive control is associated with default-executive coupling in aging (Spreng & Turner, 2019). However, without longitudinal FC measures, this study cannot say if the changes in cognitive control were accompanied by changes in dlPFC-DMN FC. Furthermore, we did only find a weak (p = 0.09) relationship in the youth group. As we had relatively fewer participant in this group with MRI scans, particularly below 20 years (n = 65), this null finding might be related to statistical power. However, a challenge when assessing Stroop abilities in children is that reading abilities are still developing which make the Stroop task less effective in capturing “pure” executive functions in children. We addressed this by correcting for color naming in addition to reading abilities in our statistical models, however, this may still be an issue, which in turn may affect how Stroop performance relates to FC in this age group. Hence, we cannot conclude that change in cognitive control only associate with dlPFC-DMNa FC during stages of aging in the lifespan and not to development and early adulthood.

Our results did not point towards specificity of FC from the left dlPFC to the DMNa in relation to loss of cognitive control. We found that left dlPFC connectivity to a network related to cognitive control processing (ControlB) was associated with a longitudinal increase in completion time, potentially demonstrating an increased demand for control processes in aging (Grady, 2012). Moreover, dlPFC connectivity with a network of external attention (DorsAttnA) was related to decreased completion time in the Stroop task. Importantly, this network profile also showed a significant negative age-slope. Hence, dlPFC-DMNa and dlPFC-DorsAttnA FC display opposite age relations and opposite relations with cognitive control change. These observations are in line with the well-established anti-correlation between networks of external attention and the DMN (Esposito et al., 2018; Fox et al., 2009). Together, our results suggest that decline in cognitive control is not specifically related to dlPFC-DMN coupling but may also associate with other dlPFC connections. This finding suggests that declining cognitive control abilities relate to broader lateral-prefrontal interconnectivity beyond just the DMN, potentially reflecting a more general pattern of dedifferentiation where functional networks become less segregated with age (Chan et al., 2014; Grady, 2012). Nevertheless, the FC profiles showing associations with change in cognitive control were all related to networks which are shown to have age-related alterations, such as, networks of cognitive control, and networks of attention (Dixon et al., 2018; Grady et al., 2016; Rieck et al., 2017). However, without longitudinal FC measures, this study cannot say that any change in cognitive control was accompanied by changes in FC.

### 4.3 Does change in level of FC relate to memory performance**?**

While resting-state FC between the dlPFC and DMN has been linked to poor memory performance in older age (Zhukovsky et al., 2022), none of the left dlPFC- network connectivity investigated here was related to memory performance during either encoding or retrieval across the lifespan after correction for multiple comparisons, except for the ControlA network. This discrepancy may be due to differences in sample characteristics; where we used a cognitively healthy lifespan sample, Zhukovsky and colleagues (2022) associated FC and episodic memory in normally aging participants over 70 years of age and compared this relationship to participants on track to get clinical dementia over 70 years of age. Although our results did not show an association between dlPFC-DMNa FC with memory performance, higher levels of FC between these two regions were observed during both encoding and retrieval with age which indicates that this this connectivity profile intrinsically relates to source memory processing.

### 4.5 Considerations of Functional Connectivity Estimation and Preprocessing

In this study, we employed cPPI analysis to estimate task-based FC. Unlike traditional FC approaches that rely on time-series correlations during rest or task periods, cPPI incorporates both intrinsic connectivity and task-evoked modulations by computing the partial correlation between region pairs while regressing out task design and physiological confounds (Fornito et al., 2012). As such, cPPI is not exclusive to a specific cognitive phase like encoding or retrieval but rather captures a more general task-related connectivity profile, including background connectivity.

To assess the reliability of our findings, we examined functional connectivity estimates both with and without global signal regression (GSR). Although GSR remains controversial due to concerns about its potential to introduce artificial anticorrelations and in some cases the removal of meaningful neural variance (K. Murphy et al., 2009; Saad et al., 2012) prior work suggests it can also enhance network specificity by removing global noise components (Fox et al., 2009; Li et al., 2019; Power et al., 2014). We observed that age related FC during encoding between dlPFC and ControlA and became non-significant without GSR. Moreover, additional networks were significantly and positively associated with age when GSR was not used as a denoising step. This might be due to spurious anticorrelations introduced by GSR (Erdoğan et al., 2016). Furthermore, without GSR fewer (2 vs 4) dlPFC-network connections related to loss of cognitive control. However, the main findings of dlPFC-DMNa associations with age and cognitive control, was consistently observed across both preprocessing approaches.

### 4.5 Limitations

A main limitation of the current study is the use of cross-sectional functional connectivity and episodic memory data, preventing us from assessing changes over time in these features in relation to the observed longitudinal changes in cognitive control. As participants with more cognitive control decline may already have lower FC at baseline, this lack of longitudinal FC data prevents answering the question of whether decline in cognitive control precedes or co-occur with FC changes. Hence, future research should investigate simultaneous changes in both FC and cognitive control to determine if these are level differences already present in FC. The age used to split the sample (30 years) was derived by combining empirical evidence and episodic memory data. However, we acknowledge that this cutoff may not align perfectly with developmental transitions or brain maturation processes occurring during this period.

## 5 Conclusion

This study showed that dlPFC-DMNa FC varied across the lifespan, with higher levels of coupling from age 30 to older adulthood and lower coupling during childhood, adolescence, and young adulthood. This dlPFC-DMNa coupling was associated with cognitive control decline in older age. Additionally, other dlPFC-to- network FC profiles also exhibited age-related changes and were linked to cognitive control loss, indicating that dlPFC-DMNa coupling is accompanied by other age- related dlPFC couplings which also are associated with cognitive decline. While this coupling was not related to episodic memory performance, it reflects age-related engagement during episodic memory processing. Future research should explore how cognitive control changes relate to longitudinal changes in memory and the coupling between dlPFC and DMN, attentional and control networks.

## Data and Code Availability

The data used in this study are not publicly available due to their sensitive nature and the involvement of human subjects. The code developed for this study is also not publicly available, as it was created for internal use only and is not maintained or documented for external distribution.

## Ethics Approval and Informed Consent Statement

This study was approved by the Regional Ethical Committee of South Norway and conducted in accordance with the principles expressed in the Declaration of Helsinki. Written informed consent was obtained from all adult participants and from parents or legal guardians of participants under the age of 18. Identifying information has not been published, and participant confidentiality has been strictly maintained throughout. All procedures followed ethical guidelines, and informed consent included agreement to participate in the study and for the data to be used in anonymized research publications.

## Author Contributions

EGB: Methodology, Formal analysis, Visualization, Writing: Original Draft, Review & Editing; HG: Conceptualization, Methodology, Formal analysis, Visualization and Writing: Review & Editing; AF: Writing: Review & Editing; DVP: Writing: Review & Editing; KBW: Writing: Review & Editing; MHS: Writing: Review & Editing.

