## Supplementary Material for "Memory-Related Default-Executive Coupling Across the Lifespan and Associations with Changes in Cognitive Control"

### Section 1. Follow-up analyses and supplementary to the main analyses.

#### 1.1. Longitudinal age-time interactions for cognitive variables

Follow-up analyses were conducted for significant age-time interactions in their linear regression models over/under the median age. There were not enough observations to estimate random intercept and slope in the linear mixed model using the median age for the aging subsamples, so we increased age by one year (51 years). For the early-life subsample, random effects exceeded the number of observations in the same analysis, so we ran the model with only random intercept. The analyses revealed significant increase in completion time over median age (50 + 1 years, range 50 – 80) (b = 2.27, t = 2.29 p = 0.02) and a not a significant change in completion time under median age (30 – 50) (b. -0.48, t = -0.55 p = 0.57). Same analyses for vocabulary revealed that age above 30 years was not associated with change in vocabulary (b = 0.19, t = 0.33 p =0.73) (See **SM Figure 1**). Follow-up analyses for the early-life subsample showed that between 8 and 22 years, completion time in the Stroop task declined with age and then it stabilized up until 30 years. Positive change in vocabulary performance was significantly associated with a higher age (b = 40.92, t = 10.37 p < 0.01).

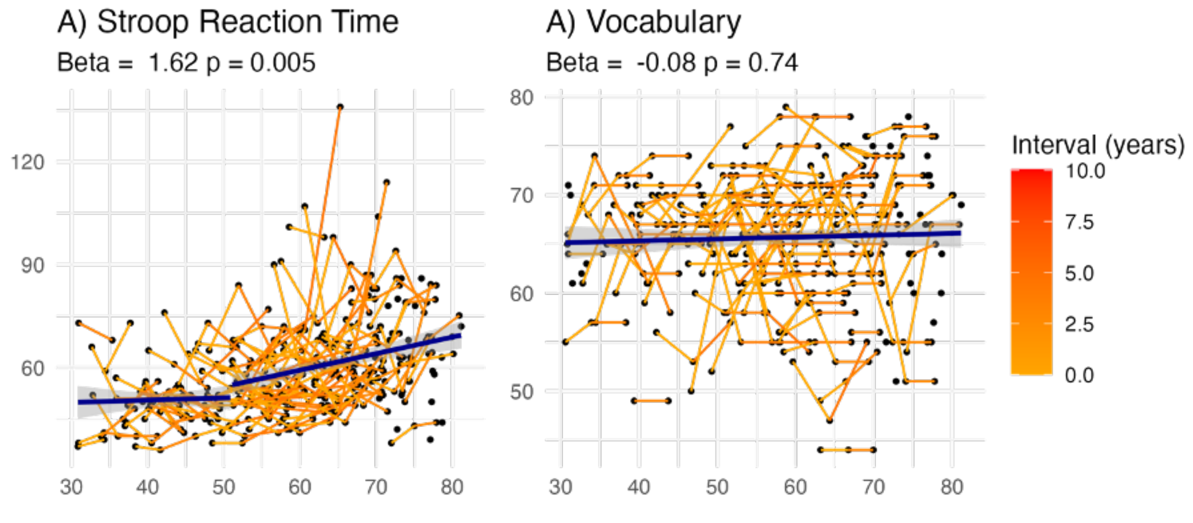

SM Figure 1. Longitudinal changes in cognitive performance for individuals aged 50 and above. The transition from orange to red signifies time in years between cognitive measurements.

##### 1.2. Memory performance varying with longitudinal Stroop performance.

We also tested for a relationship between memory performance and change in Stroop completion time. Results showed that in the aging subsample there was a linear association between memory performance and change in completion time (edf = 1.00, F. = 6.12 p = 0.01, **SM Figure 2**), namely that an increase in completion time was related to lower memory performance. In the early-life subsample, longitudinal change in completion time was not related to memory performance (edf = 1, F = 0.0, p = 0.99)

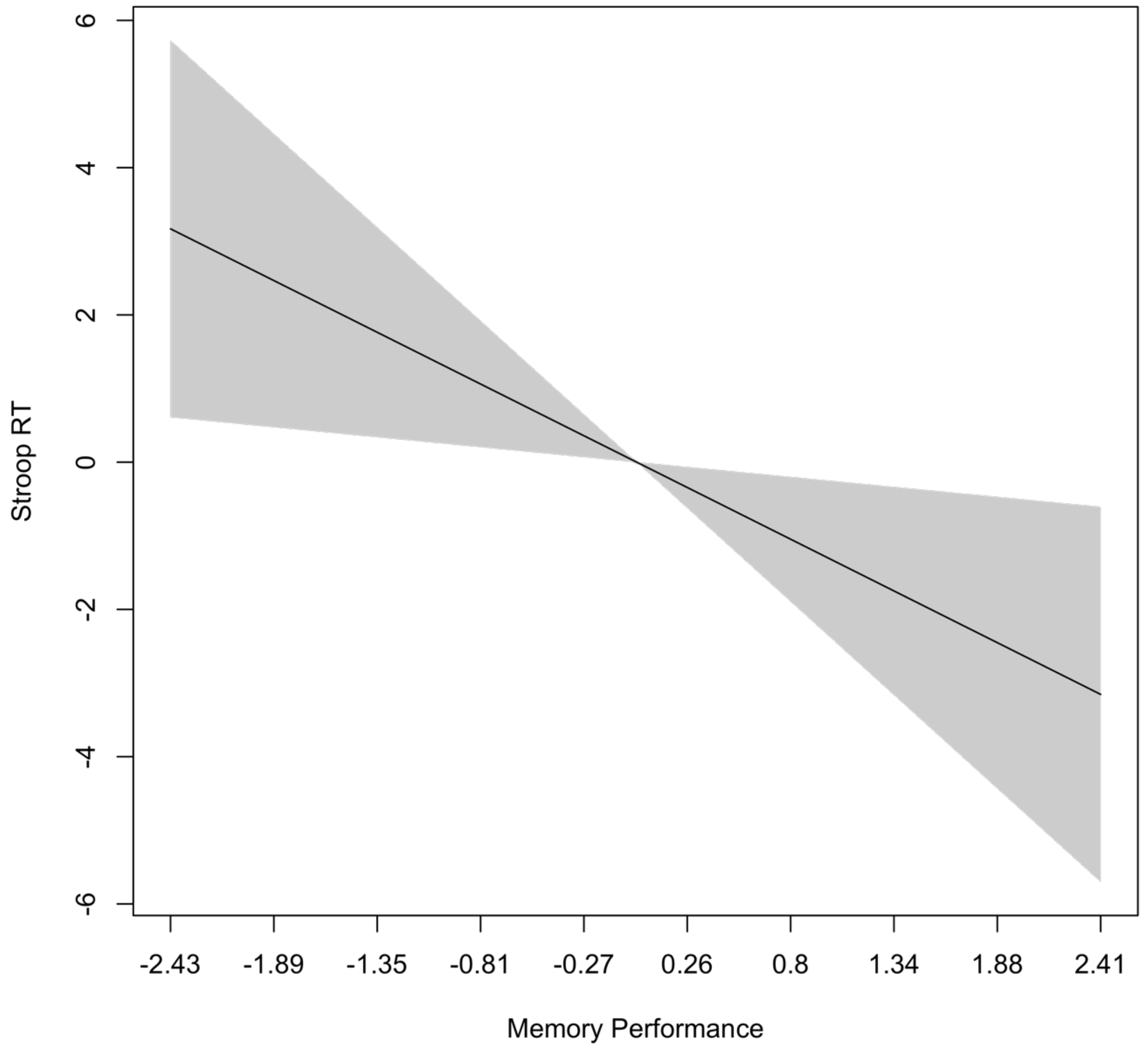

SM Figure 2. Longitudinal Stroop performance varying with memory performance. On the y-axis, Stroop completion time is represented, with positive values indicating an increase in completion time. The x-axis reflects performance in the in-scanner memory task.

#### 1.3. Age-FC associations

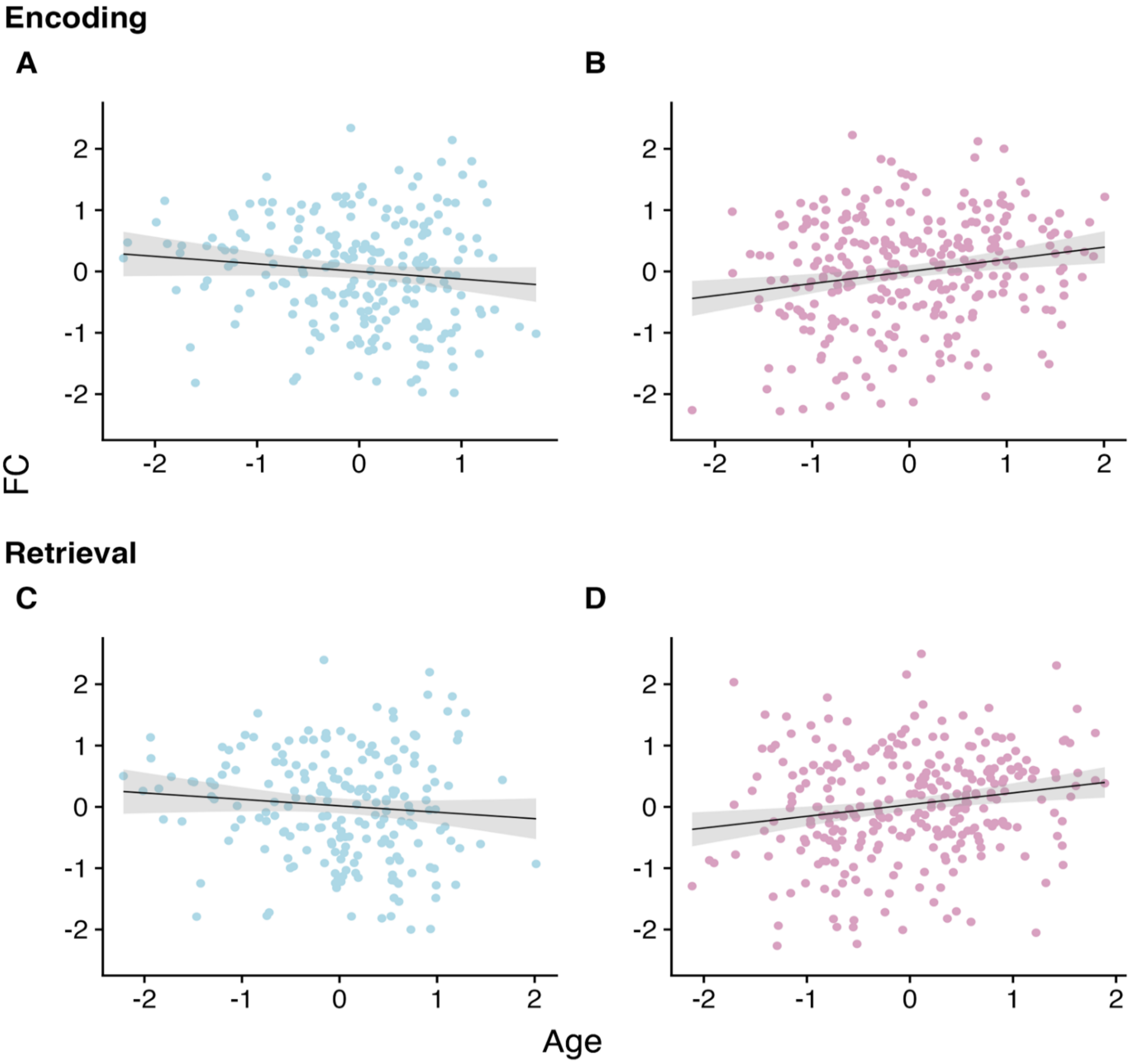

SM Figure 3. Age trajectories of dlPFC-DMNa FC divided in memory phase and age group, displaying residuals after removing sex, within-subject FC signal, and motion. **A)** Age relations for the early-life subgroup during encoding, **B)** age relations for the aging subgroup during encoding, **C)** age relations for the early-life subgroup during retrieval, and **D)** age relations for the aging subgroup during retrieval.

#### 1.4. Main effects of FC on memory performance

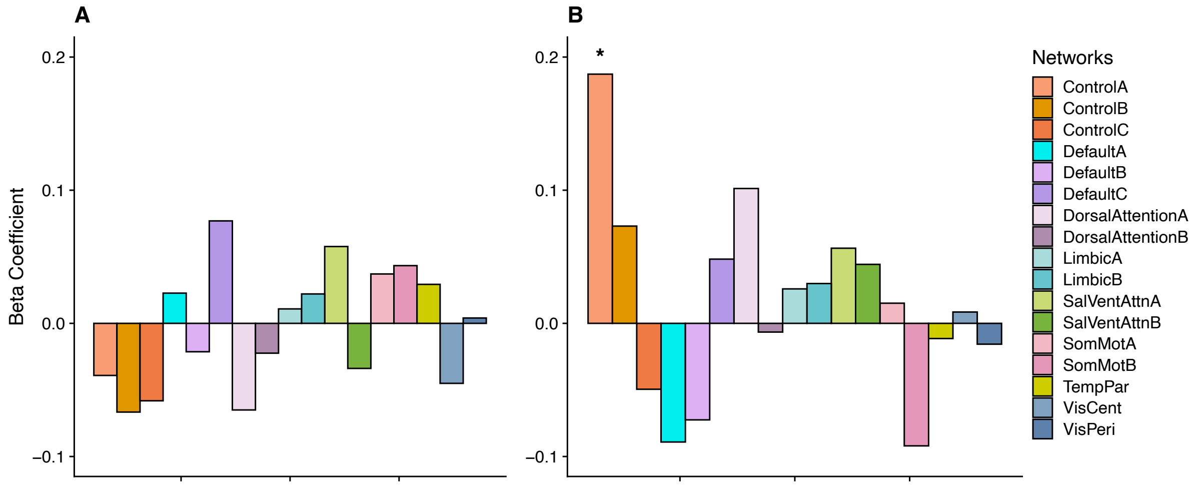

SM Figure 4. All dlPFC connections and associated memory effects, with the y-axis representing the beta coefficient values derived from linear regression models. Coefficients with P < 0.05 are indicated by an asterisk. Note that no P values survived correction for multiple comparisons (FDR). **A)** Encoding and **B)** retrieval.

#### 1.5. Age-FC interactions on memory performance

The retrieval phase showed age-FC interactions for dlPFC-LimbicB FC**,** b = -0.02, t = 2.43 p = 0.01) and SalventAttnA (b =0.02, t = 2.03 p = 0.04) before corrections for multiple comparisons. Here, follow-up analyses demonstrated that under median age (50 years, range 30 – 50 years), FC between dlPFC and LimbicB was positively associated with better memory performance (b. = 0.13, t = 2.00 p = 0.04) while over median age (range 50 – 81) was not associated with memory performance, meanwhile for SalventAttnA the slopes went in opposite direction for participants under (b =0.07, t = 0.74 p = 0.45) and over median age (b =-0.08, t = -1.61 p = 0.10), but none of the slopes were significant. See **SM Figure 5.**

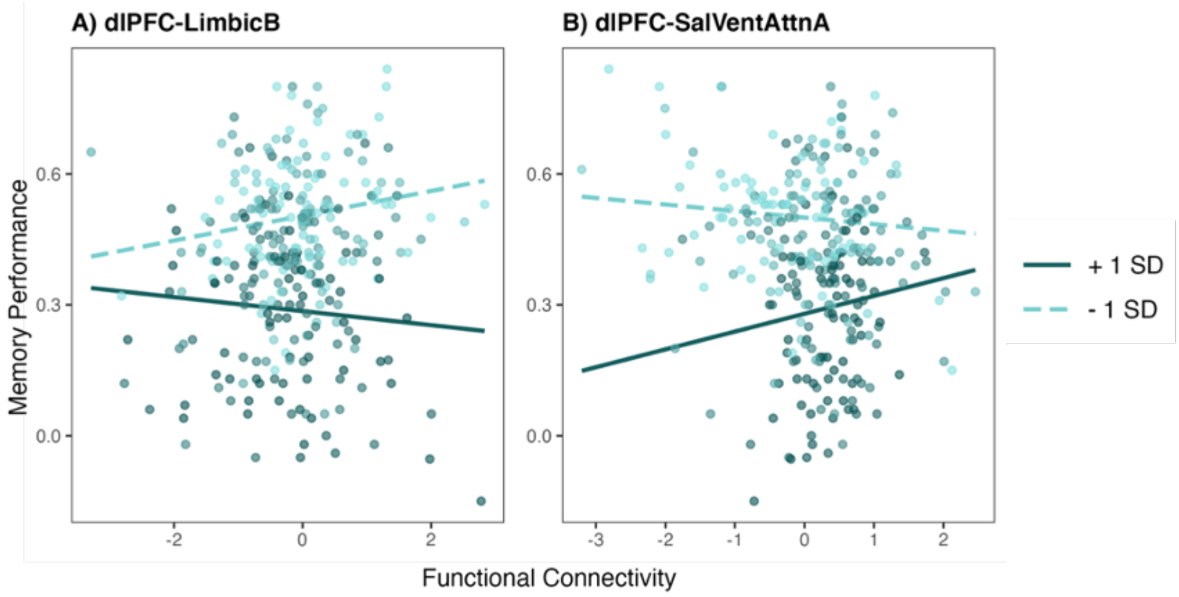

SM Figure 5. Age-FC interactions on memory performance. Lines are divided into +- 1 SD from the mean age.

### Section 2. Results without GSR as denoising step.

#### 2.1 dlPFC–DMNa Functional Connectivity and Age

We first tested the hypothesis that dlPFC–DMNa functional connectivity (FC) varies with age differently across the lifespan and memory phases. In the aging subsample, higher age was significantly associated with increased FC during both encoding ($b = 0.09$, $t = 2.30$, $p = 0.02$, FDRp = 0.03) and retrieval ($b = 0.08$, $t = 2.27$, $p = 0.02$, FDRp = 0.04). In the early-life subsample, FC was not significantly related to age during encoding ($b = -0.02$, $t = -0.69$, $p = 0.49$), but was negatively associated during retrieval ($b = -0.08$, $t = -2.40$, $p = 0.01$). All reported p-values survived correction for multiple comparisons (see **SM Figure 6**)

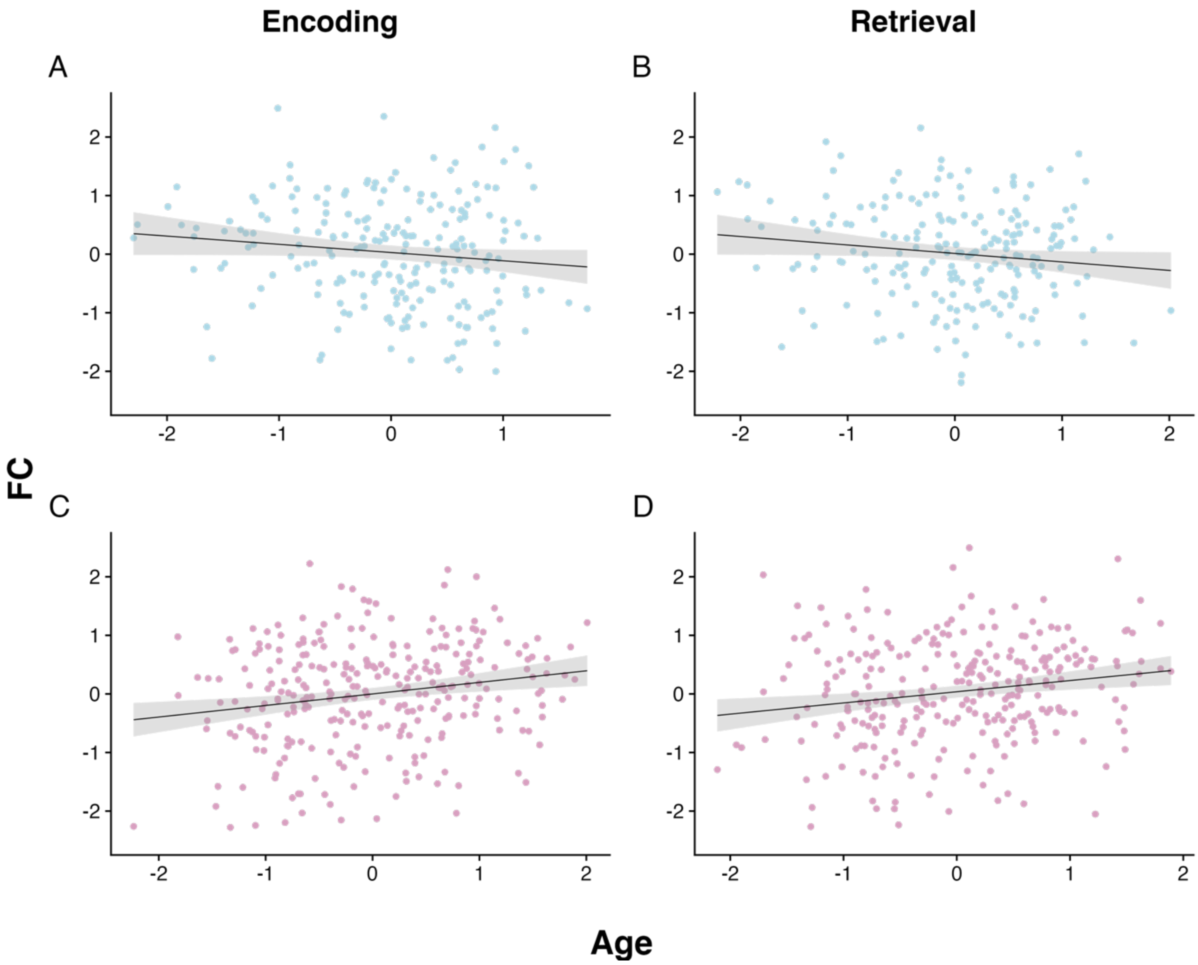

SM Figure 6. Age associations with dlPFC-DMNa FC. Displayed are model residuals in which the effects of in-scanner motion, sex, and average cortical FC have been removed from both the FC and age variables. **A)** Age trajectory of dlPFC-DMNa for the early-life subsample during encoding. **B)** Age trajectory of dlPFC-DMNa for the early-life subsample during retrieval. **C)** Age trajectory of dlPFC-DMNa for the aging subsample during encoding. **D)** Age trajectory of dlPFC-DMNa for the aging subsample during retrieval.

#### 2.2 Age-Related FC Associations in Other Networks

To assess whether age-related FC patterns were specific to dlPFC–DMNa, we examined FC between the dlPFC and 16 additional networks across memory phases (see **SM Figure 7**).

In the aging group during encoding, age was positively associated with FC between dlPFC and several networks, including ContC (b = 0.26, t = 6.43, p < 0.001), SalVentAttnA (b = 0.11, t = 2.94, p = 0.003), SomMotA (b = 0.16, t = 3.99, p < 0.001), SomMotB (b = 0.08, t = 2.15, p = 0.03), DorsAttnA (b = 0.13, t = 2.88, p = 0.004), DorsAttnB (b = 0.16, t = 3.81, p < 0.001), SalVentAttnB (b = 0.10, t = 2.62, p = 0.009), TempPar (b = 0.09, t = 2.25, p = 0.01), VisCent (b = 0.21, t = 4.27, p < 0.001), and VisPeri (b = 0.10, t = 2.32, p = 0.02).

During retrieval, the aging group showed positive associations between age and FC for SalVentAttnA (b = 0.09, t = 2.64, p = 0.008), SomMotA (b = 0.12, t = 3.47, p < 0.001), SomMotB (b = 0.11, t = 3.01, p = 0.02), DorsAttnA (b = 0.10, t = 2.55, p = 0.01), DorsAttnB (b = 0.12, t = 3.40, p < 0.001), , TempPar (b = 0.09, t = 2.50, p = 0.01), VisCent (b = 0.21, t = 4.47, p < 0.001), and VisPeri (b = 0.13, t = 2.86, p = 0.004),. A negative age slope was observed for ContA (b = -0.11, t = -2.48, p = 0.004). All findings in the aging group survived correction for multiple comparisons.

In contrast, the early-life subsample showed fewer and less robust associations. During encoding, only a negative age slope with DMNc survived correction (b = -0.12, t = -3.09, p = 0.002). During retrieval, nominal associations were observed for VisCent (b = -0.10, t = 2.16, p = 0.03) and LimbicB (b = -0.11, t = -2.55, p = 0.01), though these did not survive multiple comparisons correction.

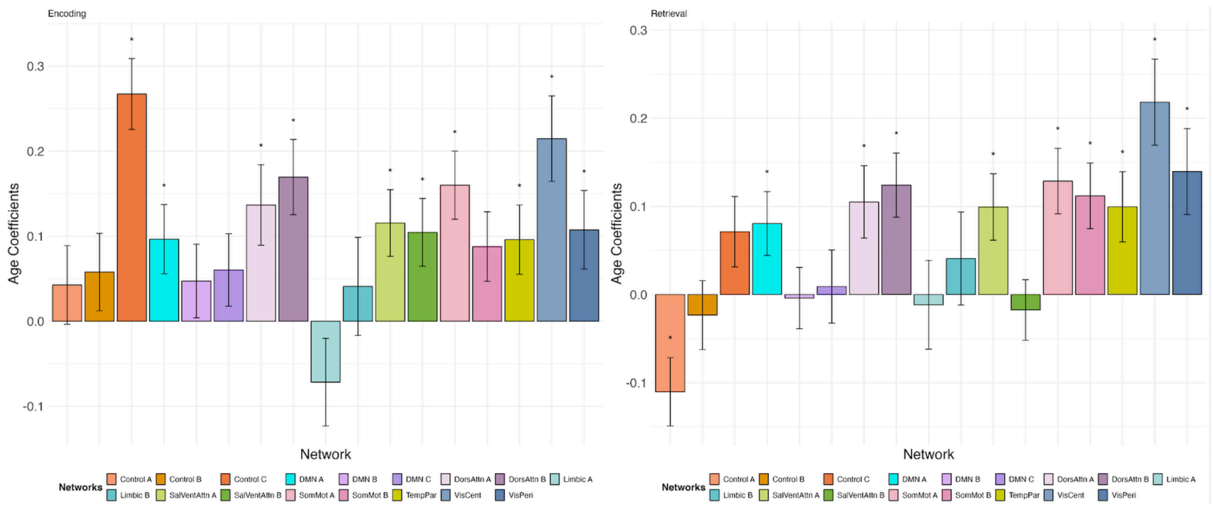

SM Figure 7. All significant age associations during encoding and retrieval for the aging subsample without GSR.

#### 2.3 Longitudinal Associations Between FC and Change in Cognitive Control

We next examined whether functional connectivity predicted longitudinal change in cognitive control, measured by Stroop inhibition completion time. In the aging subgroup during encoding, higher dlPFC–DMNa FC was associated with greater increases in Stroop completion time (edf = 1.00, F = 5.24, p = 0.02), suggesting decreased cognitive control over time (**SM Figure 8**).

During retrieval in the aging group, a significant association was found for FC between dlPFC and DorsAttnB, where higher FC predicted changes in Stroop completion time (edf = 2.90, F = 3.46, p = 0.01). No significant FC–cognitive control associations were observed in the early-life group during either encoding or retrieval.

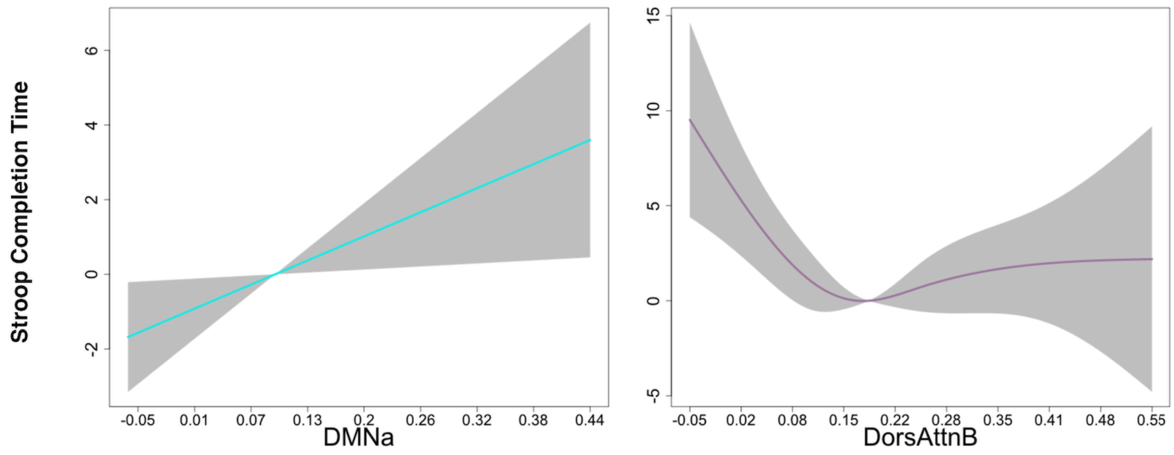

SM Figure 8. Longitudinal change in cognitive control, as measured by Stroop completion time, varying with dlPFC-to-network FC. On the y-axis, change in completion time is represented, with positive values indicating an increase in completion time. The x-axis reflects the level of functional connectivity. Both networks remained significant after correction for multiple comparisons and without GSR as denoising step.

#### 2.4 Associations Between FC and Memory Performance

Finally, we tested whether functional connectivity during memory tasks was associated with memory performance. In the aging group, no significant associations were found between memory performance and FC between dlPFC and DMNa during encoding ($b = 0.07$, $t = 0.81$, $p = 0.41$) or retrieval ($b = -0.13$, $t = -1.35$, $p = 0.18$). No other networks showed significant associations with memory performance. Similarly, in the early-life group, no FC–memory relationships reached significance.

### Section 3. Uncorrected FC results from the main analyses (with GSR).

#### 3.1. Uncorrected age relations with FC

In addition to the corrected significant main findings, during encoding for the aging subsample, age was related to higher FC between dlPFC and VisPeri (p. = 0.03) and SomMotA (p. = 0.02). At retrieval, similar positive relations with age were obtained for VisPeri (p = 0.02). In the early-life subsample, higher age was related to lower FC between dlPFC and DMNc (p = 0.01) and SalventAttna (p = 0.02) during encoding and to higher FC for ContB (p = 0.03). During retrieval, higher age was related to lower FC between dlPFC and SomMotA (p = 0.04), and to higher FC for VisCent (0.01) and ContC (0.04), respectively. SomMotA showed an opposite relation with higher age in the early-life subsample compared with the aging subsample.

#### 3.2. Uncorrected Stroop-FC associations

For the aging subsample during encoding, dlPFC-DMNb (edf = 1.00, 4.55 p = 0.03), and dlPFC-DMNc (edf = 3.22, F = 2.61 p = 0.02) were significantly associated with change in stroop completion time before correction for multiple comparisons. At retrieval, dlPFC-VisPeri FC had a positive association with Stroop completion time (edf = 3.32, F = 4.61 p = 0.01) (See **SM Figure 9)**.

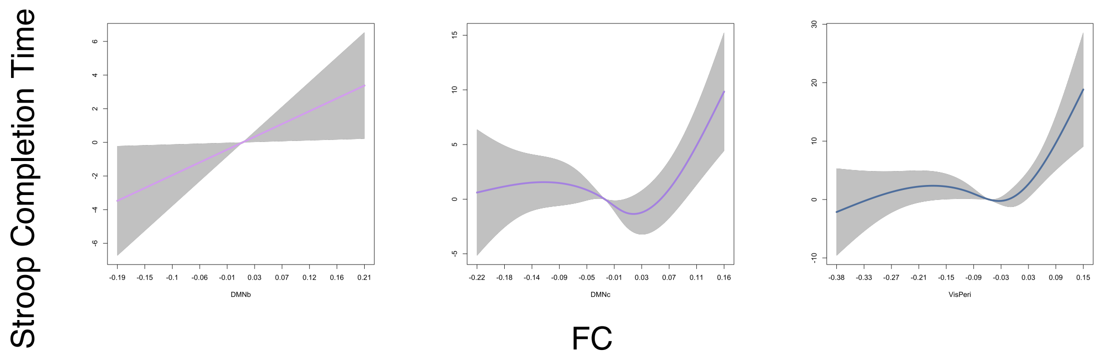

SM Figure 9**.** Uncorrected longitudinal change in cognitive control, as measured by Stroop completion time, varying with dlPFC-network FC during encoding in the aging subgroup. On the y-axis, change in completion time is represented, with positive values indicating an increase in completion time. The x-axis reflects the level of functional connectivity. **Left panel** = DMNb FC during encoding. **Middle panel** = DMNc FC during encoding. **Right panel** = Visperi during retrieval.

For the early-life subsample FC increased between dlPFC and ContA (edf = 1.00, F = 4.09 p = 0.04), ContB (edf =1.00, F = 4.13 p = 0.04), and VisPeri (edf = 1, F = 4.18 p = 0.03), and, during retrieval, ContA (edf = 3.36, F = 4.86 p = 0.003) also increased. dlPFC-ContB was the only significant network in both age groups with similar change trajectories in Stroop completion time (See **SM Figure 10)**

**SM Figure 10**

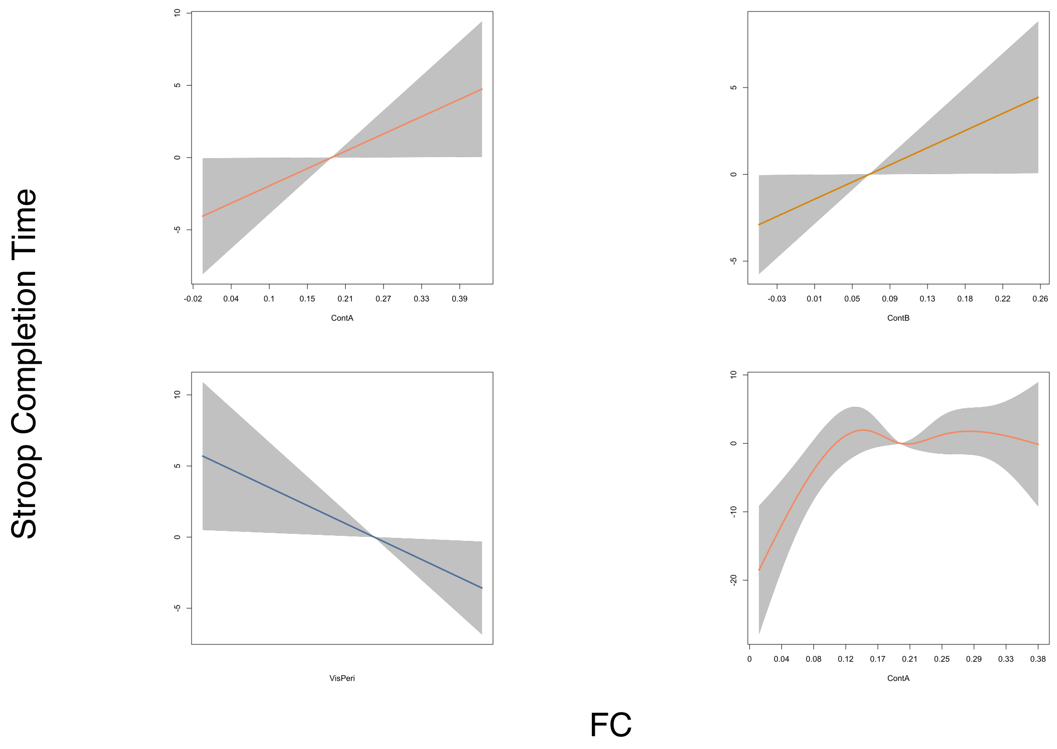

SM Figure 10. Uncorrected longitudinal change in cognitive control, as measured by Stroop completion time, varying with dlPFC-network FC during encoding in the early-life subgroup. On the y-axis, change in completion time is represented, with positive values indicating an increase in completion time. The x-axis reflects the level of functional connectivity. The upper row illustrates the aging subsample: **Upper left panel** = ContA FC during encoding. **Upper right panel** = ContA FC during retrieval. **Lower right panel** = VisPeri FC during encoding. **Lower left panel** = ContB during encoding.
